# Major threats to a migratory raptor vary geographically along the eastern Mediterranean flyway

**DOI:** 10.1101/2020.12.16.422983

**Authors:** Steffen Oppel, Volen Arkumarev, Samuel Bakari, Vladimir Dobrev, Victoria Saravia, Solomon Adefolu, Lale Aktay Sözüer, Paul Tersoo Apeverga, Şafak Arslan, Yahkat Barshep, Taulant Bino, Anastasios Bounas, Turan Çetin, Maher Dayyoub, Dobromir Dobrev, Klea Duro, Laith El-Moghrabi, Hana ElSafoury, Ahmed Endris, Nabegh Ghazal Asswad, Junior Hanson Harry, Sam T Ivande, Sharif Jbour, Eleftherios Kapsalis, Elzbieta Kret, Bruktawit A Mahamued, Shiiwua A Manu, Solomon Mengistu, Abdoul Razack Moussa Zabeirou, Sulaiman Inuwa Muhammad, Slave Nakev, Alex Ngari, Joseph Onoja, Maher Osta, Serdar Özuslu, Nenad Petrovski, Georgi Popgeorgiev, Cloé Pourchier, Alazar Ruffo, Mohammed Shobrak, Lavrentis Sidiropoulos, Theodora Skartsi, Özgün Sözüer, Kalliopi Stara, Million Tesfaye, Mirjan Topi, Dimitrios Vavylis, Metodija Velevski, Zydjon Vorpsi, Mengistu Wondafrash, Erald Xeka, Can Yeniyurt, Emil Yordanov, Stoyan C Nikolov

## Abstract

Millions of large soaring birds migrate from the Palaearctic to Africa every year, and follow distinct flyways around the Mediterranean Sea. While there is conservation concern for many long-distance migratory bird populations, the magnitude and geographic range of threats affecting birds along flyways are poorly known, which complicates efficient mitigation. We used an endangered soaring migrant, the Egyptian Vulture *Neophron percnopterus*, as an example species to assess important threats in 13 countries along the eastern Mediterranean flyway. We tracked 71 birds using satellite telemetry to quantify mortalities, surveyed 4198 km of powerlines to detect dead birds, conducted 910 interviews to quantify the prevalence of poison use, and assessed the magnitude of direct persecution by surveying markets and hunters. We lost 44 birds (50% in Europe and the Mediterranean Sea, 16% in the Middle East, and 34% in Africa), and mortality causes varied geographically. Inadvertent poisoning resulting from rural stakeholders targeting predators occurred along most of the flyway. On the breeding grounds in eastern Europe, poisoning and collision and electrocution continue to be major threats. Electrocution on small and poorly designed electricity pylons was most severe in Turkey, Ethiopia and Saudi Arabia, while direct persecution to meet market demands for belief-based use of vulture products appears to be the largest threat in Nigeria and Niger. Illegal direct persecution for leisure is a major threat in the Middle East and Egypt. Although our work cannot quantitatively estimate which of the identified threats has the greatest demographic impact on Egyptian Vultures, none of threats are species-specific and are therefore relevant for many other migratory birds. Our assessment highlights the key threats per country that range states need to address to meet their obligations under the Convention of Migratory Species to protect migratory birds.

## Introduction

Migratory animals connect countries and continents, and pose a particular conservation challenge, because threats to migratory animals that occur in one geographic region may limit their population size at distant breeding or wintering regions. The global network of protected areas, established to conserve biodiversity and habitats, is however largely inadequate to protect migratory animals along their flyways (Runge *et al*. 2015). Countries through which migratory animals pass have therefore committed to internationally coordinated conservation measures to protect migratory animals throughout their range under the Convention of Migratory Species, an environmental treaty of the United Nations.

Every year millions of large soaring birds migrate from European breeding grounds to wintering areas in Africa (Porter & Beaman 1985). Because most soaring birds are reluctant to cross large water bodies such as the Mediterranean Sea (Agostini *et al*. 2015), major flyways exist that follow routes around either the western or eastern periphery of the Mediterranean Sea (Porter & Willis 1968, Finlayson 1992, Shirihai *et al*. 2000). In the east, a major flyway funnels birds from Central and Eastern Europe and western Asia around the Black Sea, via the Bosporus in Turkey, and the Middle East towards the Red Sea, which many birds either pass to the north near Suez in Egypt or cross in the south at Bab-el-Mandeb between Yemen and Djibouti (Bijlsma 1983, Welch & Welch 1988, Phipps *et al*. 2019).

Counts of migrating raptors along this flyway indicate that >1 million raptors of >25 species regularly migrate along this route (Alon *et al*. 2004, Verhelst *et al*. 2011, Fülöp *et al*. 2014). Among these species are 12 globally threatened species, and several whose populations appear to be declining on breeding grounds (Brochet *et al*. 2016). Because populations of migratory birds can be limited by events that occur outside their breeding areas, understanding the threats along the flyway is important to inform effective conservation. Although many of the threats that exist along the flyway are broadly known in general terms (Kirby *et al*. 2008, Ogada *et al*. 2015, Brochet *et al*. 2016), there is so far no detailed analysis describing which threats pose the greatest risk in which geographic region to efficiently target conservation efforts.

One of the globally threatened species using the eastern Mediterranean flyway is the Egyptian Vulture *Neophron percnopterus*, a typical soaring migrant with a broadly dispersed wintering range (Buechley *et al*. 2018, Phipps *et al*. 2019) representative of many other species along the flyway (Porter & Beaman 1985). The Egyptian Vulture breeding population in the Balkans in eastern Europe has declined dramatically over the past 30 years (Velevski *et al*. 2015, Arkumarev *et al*. 2018), and while threats on breeding grounds have been studied and partially mitigated, the magnitude and distribution of threats along the flyway is so far poorly understood.

Here we examine the relative intensity of different known major threats to Egyptian Vultures in 13 countries along the eastern Mediterranean flyway, and present a qualitative assessment of their relative importance. We first collected quantitative evidence of bird mortality through satellite tracking, powerline surveys and interviews with local people to assess poisoning and persecution. We then used this quantitative evidence in an expert consultation to qualitatively rank the most important threats by geographic region. This country-specific threat ranking can be used by range states to prioritise conservation work to meet their obligations under the Convention of Migratory Species.

## Methods

### Study species and study region

The Balkan population of Egyptian Vultures breeds in Albania, North Macedonia, Bulgaria, Turkey and Greece (Velevski *et al*. 2015), and migrates to wintering areas in Africa and the southern Arabian peninsula (Oppel *et al*. 2015, Buechley *et al*. 2018, Phipps *et al*. 2019). We focussed our study on the five breeding countries, and important countries along the flyway and wintering areas where work was practically feasible. Specifically, we conducted work in Albania, North Macedonia, Bulgaria, Greece, Turkey, Syria, Lebanon, Jordan, Egypt, Saudi Arabia, Ethiopia, Niger and Nigeria. All these countries, except Turkey, are parties of the Convention of Migratory Species and have therefore committed to protecting migratory animals. Study areas within countries were selected based on knowledge about the distribution of birds from surveys (Hilgerloh *et al*. 2011, Arkumarev *et al*. 2014, Oppel *et al*. 2014) or from satellite telemetry (Oppel *et al*. 2015, Buechley *et al*. 2018).

### Telemetry to assess causes of mortality and collision risk with wind turbines

From August 2010 to October 2020 we tracked 71 individually marked Egyptian Vultures (46 juveniles, 16 immatures, 9 adults) with solar-powered GPS transmitters (Microwave or Ornitela, 30-45g, all <3% of body mass) from capture locations in the Balkans, Ethiopia, and Jordan at high spatial and temporal (one GPS position every 10 min – 1 hr) resolution (for more details see Oppel *et al*. 2015, Buechley *et al*. 2018, Phipps *et al*. 2019). Transmitters were attached with a backpack or leg-loop harness configuration that is unlikely to affect survival probability in vultures (Sergio *et al*. 2015, Anderson *et al*. 2020). For those birds whose transmissions ended, we attempted to identify the cause of signal loss through inspection of the last signals (Sergio *et al*. 2019) and ground searches. When carcasses were found or the cause of signal loss could be reliably ascertained from the sequence of transmissions, we classified mortality as either ‘natural mortality’ (e.g. drowning, predation, exhaustion, starvation), ‘direct persecution’, ‘poisoning’ or ‘electrocution’; all remaining cases were classified as ‘unknown’. We present proportions of the number of tracked animals and of identifiable mortalities that were due to different causes.

Collision with wind turbines is a potential mortality cause for Egyptian Vultures (Carrete *et al*. 2009, Thaxter *et al*. 2017), but we did not observe mortality from wind turbine collision in our tracking data. We therefore used these tracking data to assess the potential exposure of birds to wind turbines along the flyway. We used data hosted on Open Street Map that summarised the location and size of wind power generation facilities (hereafter ‘windfarms’) (Dunnett *et al*. 2020), supplemented by a proprietary data set provided by The WindPower.net (obtained in June 2020). Given the spatial and temporal resolution of our tracking data, and the poor precision of turbine locations across the cross-continental scale of our study, it was impossible to assess how often a bird flew within the rotor-swept area of an existing turbine. We therefore assessed the ‘potential exposure’ to windfarm mortality by quantifying the frequency of locations within a 10 km radius around the central point of each windfarm location. A 15 km radius around wind turbines was previously found to predict mortality in Egyptian Vultures (Carrete *et al*. 2009), but we curtailed this radius to 10 km given that a tracked bird flying at 40 km/h would register at least one GPS location every 15 min within that radius.

We interpolated our tracking data to regular 15 min intervals, and summarised the total amount of time our tracked birds spent within the 10 km buffers of known windfarms. We then related the amount of time spent in the vicinity of windfarms to the total amount of time our tracked birds spent in each country to assess the relative exposure to windfarm collision risk. We acknowledge that this coarse assessment of potential exposure cannot quantify the immediate mortality risk at every single turbine, which would be affected by the turbine height, the flight altitude of the bird, and the exact flight trajectory (Marques *et al*. 2014).

### Surveys to detect electrocution and collision victims

Electrocution and powerline collision mortality is a recognised major threat along the flyway (AEWA 2012, Angelov *et al*. 2013). We therefore conducted surveys under power distribution lines in areas that were frequented by Egyptian Vultures during breeding, migration, or wintering (Fig. 1). Power lines were selected as those that were perceived to pose a high risk due to location and design of the support structure (Lehman *et al*. 2007, Bernardino *et al*. 2018, D’Amico *et al*. 2019). We selected small-to medium-voltage distribution lines that were supported by single poles with a cross-bar and conducting wires propped up above the support structure, as these powerlines are the most dangerous for electrocution (Lehman *et al*. 2007, Eccleston & Harness 2018). We spatially selected lines that were in the vicinity of breeding or feeding areas such as vulture restaurants, abattoirs or rubbish dumps, or major concentration areas on migratory routes where birds may be vulnerable when entering or departing from overnight roosts. Due to the non-random selection of searched powerlines and the heterogenous distribution of mortality (see Results), we are unable to extrapolate the total number of electrocution or collision victims per country based on the size of the power distribution network.

**Fig. 1.**
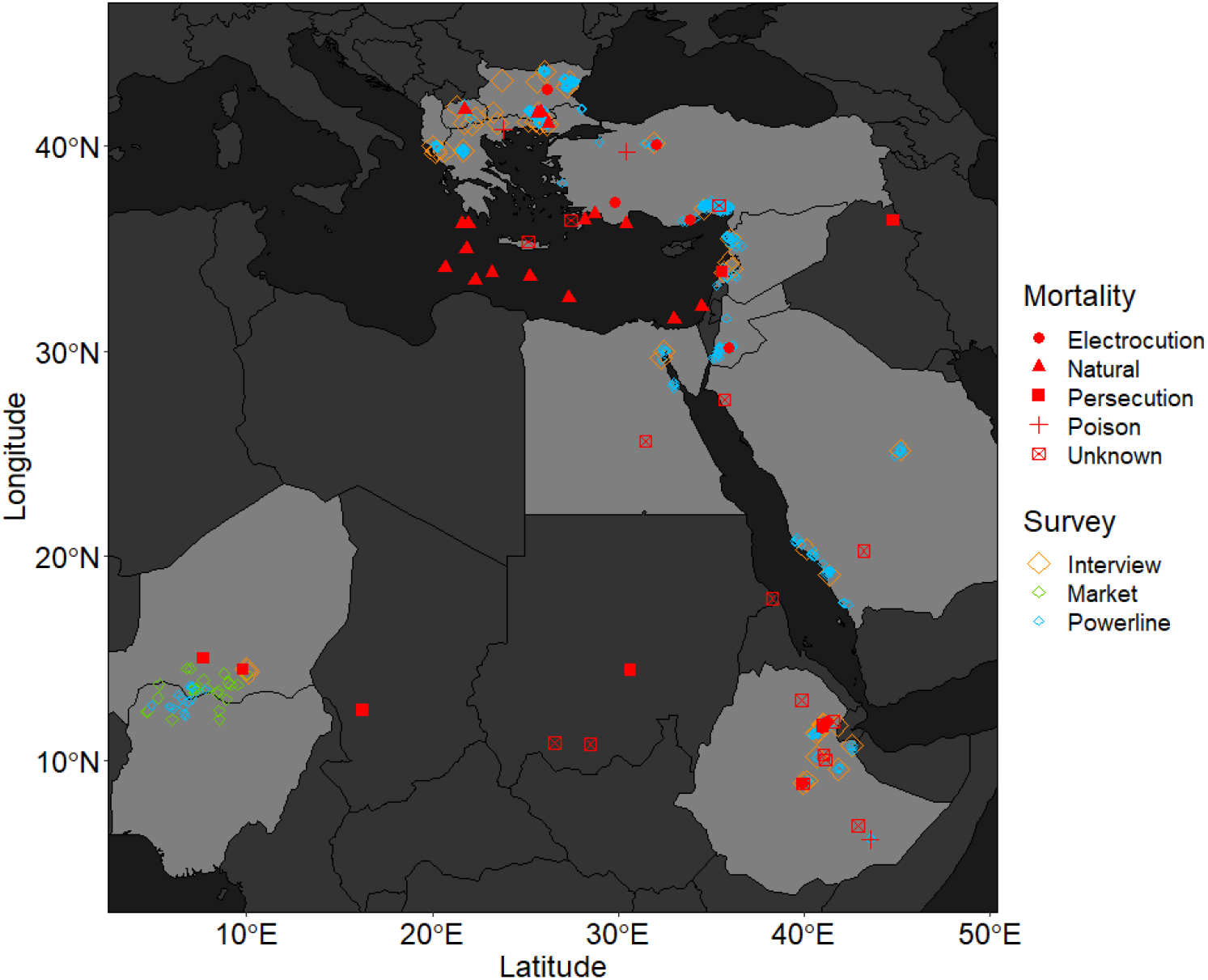
Locations of Egyptian Vulture mortality and overview of field survey effort in 13 focal countries (light grey) along the Eastern Mediterranean flyway between 2010 and 2020. Red symbols indicate locations where Egyptian Vulture mortality was recorded from either satellite-tracked individuals or from surveys.

Surveys were conducted in 2012 and 2013 (Bulgaria, Greece), and between 2018 and 2020 (remaining countries) by slowly walking under each power line and searching the ground and any adjacent vegetation where scavengers may have dragged collision or electrocution victims (Costantini *et al*. 2017). Any carcass or remnants of a carcass were identified to species, and the cause of death was assessed as electrocution or collision based on the location of the carcass and any recognisable injuries (Demerdzhiev 2014). Species and mortality causes that could not be identified were recorded as ‘unknown’. Because high mortality observed during a survey could have in theory been caused by chance if the accident happened immediately prior to the survey, we re-surveyed two lines in Ethiopia and six lines in Egypt repeatedly to exclude that the mortality found under those lines during the first survey had occurred solely by chance. Carcasses found during a survey were removed or labelled to avoid counting the same carcass again on a subsequent survey.

We summarise and present results as the relative number of victims per km of powerline surveyed in each country, but caution that this will be a minimum figure as various factors that influence detection and persistence probability of carcasses are not accounted for by our method (Bellan *et al*. 2013, Etterson 2013, Korner-Nievergelt *et al*. 2015). However, despite the crude summary, these surveys are nonetheless valuable to prioritise whether electrocution and collision are a relevant threat in a specific region and can therefore be used for prioritisation of conservation actions (Bernardino *et al*. 2018, D’Amico *et al*. 2019).

### Interviews to assess risk of poisoning

The inadvertent poisoning of vultures as a consequence of human-wildlife and human-human conflicts is a major threat on breeding (Margalida *et al*. 2014, Sanz-Aguilar *et al*. 2015, Ntemiri *et al*. 2018, Parvanov *et al*. 2018) and on wintering grounds (Ogada *et al*. 2015, Monadjem *et al*. 2018, Murn & Botha 2018). To assess the prevalence of poison use by people using rural habitat in vulture breeding or wintering areas, we conducted interviews with rural stakeholders such as livestock owners, farmers, hunters, or other people who may have conflicts with wildlife or other land users that could conceivably be resolved with poison (Santangeli *et al*. 2016, Craig *et al*. 2018). We conducted interviews in all focal countries except Jordan and Nigeria, where poisoning by rural people was considered unlikely to affect Egyptian Vultures.

Interviews were conducted in a semi-structured conversation, and in each interview rural people were asked only two general questions: (1) what are your main problems in raising livestock or growing crops?; and (2) what solutions do you routinely employ to overcome these problems? We did not ask specifically for the use of poison, and thus avoided directly asking for illegitimate behaviour that would have required sophisticated elicitation techniques (Nuno & St. John 2015, Hinsley *et al*. 2019), which would not be comparable across the cultural diversity of our study area. Answers were categorised into six different themes to portray problems (habitat availability, climate, invasive species, disease, predation, other), and into eight different types of solutions (re-location, guarding/chasing, fencing, shooting, trapping, poisoning, medicine, and other solutions). Depending on the availability and social structure of rural communities, questionnaires were conducted either with individuals or with focus groups consisting of 2-12 people. We summarised the frequency of answers per interview regardless of the number of people participating in a given interview, because group interview participants provided a single collective answer and not independent answers.

All interviews were conducted anonymously (no identity-revealing information was requested from or provided by participants) in the local language of rural stakeholders by a local associate who then translated responses back to researchers (for general locations see Fig. 1). All participants were informed of the research prior to the interview commencing, and all participants were given the opportunity to discontinue the conversation at any time (St John *et al*. 2012, St. John *et al*. 2014).

Poisoning of vultures also occurs through the use of certain veterinary medical products that can cause kidney failure in vultures (Oaks *et al*. 2004, Shultz *et al*. 2004, Green *et al*. 2006), or through the use of highly toxic substances to control agricultural pests or feral animals near human settlements (Abebe 2013, Ogada 2014, Parvanov *et al*. 2018, Plaza *et al*. 2019). We therefore interviewed officials from local authorities to find out what veterinary medical products were authorised for use in each country, and interviewed public health authorities to find out whether public health issues were controlled in ways that could put scavengers at risk of poisoning (Zewdu *et al*. 2010, Tschopp *et al*. 2016). This information was not used in a quantitative way, but merely to understand whether pharmaceuticals and toxins that have caused vulture population collapses elsewhere are in widespread use along the flyway.

### Assessment of direct persecution

Many migratory birds are directly persecuted and illegally killed or taken on migration along the entire flyway (Brochet *et al*. 2016). In addition, direct illegal persecution of vultures occurs in many African countries due to market demands of vulture products for belief-based use (Saidu & Buij 2013, Ogada *et al*. 2015, Buij *et al*. 2016). We aimed to assess the magnitude of these two processes on Egyptian Vultures by extrapolating the number of birds illegally killed through direct persecution (Brochet *et al*. 2016, Brochet *et al*. 2019a, Brochet *et al*. 2019b), and by conducting surveys of markets in Africa to assess the magnitude of the threat of market demand on vulture populations specifically. We assume that our surveys are indicative of general patterns in the countries we surveyed, but acknowledge that due to the illegal nature of direct persecution our surveys could potentially underestimate the true scale of this threat.

To assess illegal killing of migratory birds along the flyway, we used data collected in comprehensive regional assessments (Brochet *et al*. 2016, Brochet *et al*. 2019a, Brochet *et al*. 2019b). Briefly, national experts and organisations were consulted between 2014 and 2018 to assess if wild birds were known or likely to be illegally killed in non-trivial numbers in their country. Sites where hunting and shooting activities were known to have been practised were surveyed to obtain information from hunters and other informants about the species being hunted and to estimate the approximate number of birds being killed at focal hunting areas. Literature and unpublished reports were consulted to gauge the extent of illegal killing, and data were extrapolated and rounded to the nearest thousand to avoid spurious precision (Brochet *et al*. 2016, Brochet *et al*. 2019a, Brochet *et al*. 2019b). We extracted the minimum and maximum of these estimates for all birds, and for Egyptian Vultures specifically.

The available regional assessments did not include our focal countries in sub-Saharan Africa (Ethiopia, Niger, Nigeria). In Africa, the illegal killing and taking of vultures occurs to satisfy market demands primarily in Nigeria. Following the death of a tracked individual Egyptian Vulture that was shot in Niger to supply a market in Nigeria (Kret *et al*. 2018), we conducted surveys on several markets in Nigeria and Niger (Fig. 1) in 2018 and 2019 to determine the magnitude of the vulture trade with a particular focus on Egyptian Vultures. During the surveys, a local native research associate first assessed how many stalls at each market sold any vulture parts, and then quantified the number of stalls that either have in the past or would sell Egyptian Vulture parts in the future. Sellers were asked about the origin of vulture parts on sale. Vulture products are not routinely sold on markets in Europe, the Middle East, Egypt or Ethiopia, hence we did not conduct market surveys in those countries.

### Qualitative ranking of threats for each country

Our work cannot quantitatively estimate which of the identified threats has the greatest demographic impact on Egyptian Vultures. We measured each threat on a different scale and in different units, and there is no realistic approach to convert our quantitative data into a common currency. To facilitate decision making in each country which threat to prioritise, we therefore convened regional experts and ranked threats based on the quantitative evidence and expert’s knowledge about their respective countries (Martin *et al*. 2012, Hugé & Mukherjee 2018). We convened experts at regional scales for Europe (North Macedonia, Albania, Bulgaria, Greece), the Middle East (Turkey, Lebanon, Syria, Jordan, Egypt), and Africa (Ethiopia, Egypt, Niger, Nigeria) to ensure regional consistency in the ranking of threats. For each threat, a country-specific ranking was obtained by considering two dimensions, namely how widespread the threat is in a respective country, and how severe the threat is in terms of causing mortality. The importance of threats was then scored qualitatively from 0 (threat absent or irrelevant) to 5 (ubiquitous and severe), and consensus among regional experts was sought through discussions and supportive evidence. Due to the qualitative nature of the ranking of threats, we emphasize that this ranking is intended to prioritise work at a country-level: while threats may be consistently scored within continents, threat rankings between continents may not be comparable. Nonetheless, this overview of the relative importance of threats allows governments and conservation organisations in each country to prioritise efforts to reduce the most important threats in each country, a fundamental advance in the conservation strategy for migratory birds (Vickery *et al*. 2014).

## Results

We lost 44 (62%; 32 marked as juveniles, 7 immatures, 5 adults) of the 71 individual Egyptian Vultures tracked with satellite telemetry by the end of October 2020, and for 15 (34%) of these birds the cause of mortality could not be ascertained. Of the 44 lost birds, 22 (50%) perished on or near the breeding grounds in eastern Europe (including 12 juveniles drowning in the Mediterranean Sea during their first autumn migration), seven (16%) in the Middle East and Arabian Peninsula, and 15 (34%) in Africa. Of the 29 individuals whose cause of mortality could be ascertained, six (21%) were shot, two (7%) were poisoned, and two (7%) were electrocuted, while the remaining 19 (65%) died from natural causes (Fig. 1).

### Electrocution and collision

We conducted 661 transect surveys in 13 countries to search for electrocution and collision victims under powerlines, covering a total of 4198 km. During these surveys we found a total of 696 bird carcasses of 54 species, of which 16 were Egyptian Vultures. The highest encounter rate of bird carcasses per km of powerline occurred in Turkey, Ethiopia, and Saudi Arabia, and the most Egyptian Vulture carcasses were found in Ethiopia (Table 1), where the largest known congregations of this species occur (Arkumarev *et al*. 2014).

**Table 1.** Summary of power line transect surveys conducted between 2012 – 2013 (Bulgaria, Greece) and 2018 – 2020 (remaining countries) including the total number of bird electrocution and collision victims found during these surveys. Note that these surveys do not account for detection probability and carcass removal by scavengers. Countries are ordered by the carcass encounter rate.

| Country | N surveys | Total distance surveyed (km) | N of bird carcasses | N of Egyptian Vulture carcasses | Carcass encounter rate (birds/km) |
| --- | --- | --- | --- | --- | --- |
| Turkey | 144 | 247.5 | 118 | 3 | 0.477 |
| Ethiopia | 67 | 227.1 | 80 | 12 | 0.352 |
| Saudi Arabia | 52 | 195.1 | 66 | 0 | 0.338 |
| Jordan | 48 | 346.3 | 98 | 1 | 0.283 |
| Lebanon | 6 | 19.7 | 5 | 0 | 0.254 |
| Bulgaria | 151 | 578.9 | 111 | 0 | 0.199 |
| North Macedonia | 10 | 25.6 | 5 | 0 | 0.196 |
| Egypt | 86 | 1648.6 | 197 | 0 | 0.119 |
| Albania | 31 | 121.0 | 7 | 0 | 0.058 |
| Nigeria | 11 | 75.7 | 3 | 0 | 0.040 |
| Niger | 5 | 79.4 | 3 | 0 | 0.038 |
| Greece | 30 | 162.8 | 2 | 0 | 0.012 |
| Syria | 20 | 471.0 | 1 | 0 | 0.002 |

Mortality of birds under powerlines was not homogenously distributed, but occurred primarily near roosting or feeding areas. For example, in southern Turkey dozens of White Storks *Ciconia ciconia* were found under a short power distribution line adjacent to a forest that is typically used as overnight roost by migrating storks. In north-eastern Ethiopia, dozens of dead vultures were found under hazardous medium-voltage distribution lines near rubbish dumps and abattoirs where vultures frequently congregate to feed.

We tracked individual birds since 2010 for up to 8 years per individual (total 16,939 bird tracking days), resulting in 28 – 6144 bird tracking days in the selected countries along the flyway (Table 2). In two countries, Greece and Turkey, the tracked birds spent >15% of their time within 10 km of existing windfarms, and in three further countries (Jordan, Bulgaria and Lebanon) the tracked birds spent >3% of their time near windfarms (Table 2). In Ethiopia, our tracked Egyptian Vultures did not spend any time near existing windfarms as the spatial distribution of windfarms (in highlands) and Egyptian Vultures (in lowlands) differed.

**Table 2.** Summary of the potential exposure of 71 tracked Egyptian Vultures to collision risk with wind power generation facilities (‘windfarms’) between 2010 and October 2020. Birds were tracked with GPS transmitters from the Balkans and from Ethiopia, yielding the number of tracking days per country and the number of tracking days within 10 km radius of windfarms in this country. Countries are ordered by the proportion of time in windfarms.

| Country | N of windfarms | Total area of windfarms | Tracked bird days in country | Tracked bird days in windfarms | proportional time in windfarms (%) |
| --- | --- | --- | --- | --- | --- |
| Greece | 199 | 1141 | 1782 | 856 | 48.0 |
| Turkey | 295 | 2007 | 1679 | 271 | 16.2 |
| Jordan | 9 | 141 | 126 | 7 | 5.8 |
| Bulgaria | 96 | 379 | 4077 | 158 | 3.9 |
| Lebanon | 1 | 2 | 28 | 1 | 3.3 |
| North Macedonia | 2 | 13 | 105 | 1 | 0.8 |
| Egypt | 22 | 816 | 516 | 3 | 0.6 |
| Syria | 1 | 2 | 279 | 0 | 0.1 |
| Saudi Arabia | 1 | 2 | 803 | 0 | 0.0 |
| Ethiopia | 15 | 173 | 6144 | 0 | 0.0 |
| Albania | 1 | 59 | 560 | 0 | 0.0 |
| Nigeria | 1 | 29 | 103 | 0 | 0.0 |
| Niger | 0 | 0 | 737 | 0 | 0.0 |

### Poisoning

We interviewed a total of 1135 rural stakeholders during 910 distinct interviews in ten countries along the flyway where Egyptian Vultures occur regularly (Table 3). Our interviews revealed that most livestock herders experienced problems with carnivores, especially wolves, hyenas, jackals, lions, foxes, and feral dogs that occasionally attacked their livestock. The threat of carnivores to livestock was highlighted in > 65% of interviews in Albania, Niger, Syria, Turkey and Ethiopia, but only in 14% of interviews in Bulgaria and 23% in Lebanon, where the lack of grazing land was the most prevalent problem (Table 3).

**Table 3.** Summary of interviews with rural stakeholders to understand common problems with land use and livestock herding, and solutions adopted to overcome these problems. Only the most common problems and solutions are shown (see Table S1 for full list). Note that poisoning is referred to at a community rather than at an individual level, and its prevalence may therefore not be comparable to shooting or protection in some countries; ‘Habitat’ refers to the problem of insufficient grazing or crop land; ‘Protection’ refers to non-lethal livestock protection measures such as fences, bomas, guarding dogs or other predator repellents. Countries are ordered by the proportion of responses that indicated poisoning. No interview data were available for Jordan and Nigeria, where poisoning by rural people was considered unlikely to affect Egyptian Vultures.

| Country | Number of interviews | Number of participants | Problems |  | Solutions |  |  |
| --- | --- | --- | --- | --- | --- | --- | --- |
|  |  |  | Predation | Habitat | Protection | Shooting | Poison |
| Syria | 4 | 6 | 100% | 100% | 0% | 100% | 100% |
| Greece* | 367 | 367 | 47% |  | 66% | 15% | 63% |
| Saudi Arabia | 8 | 10 | 100% | 0% | 0% | 100% | 50% |
| Egypt | 4 | 4 | 50% | 25% | 25% | 0% | 50% |
| North Macedonia | 31 | 37 | 42% | 32% | 77% | 19% | 48% |
| Ethiopia | 68 | 190 | 87% | 29% | 35% | 68% | 46% |
| Niger | 6 | 46 | 67% | 50% | 83% | 0% | 33% |
| Lebanon | 22 | 53 | 23% | 45% | 14% | 32% | 27% |
| Albania | 148 | 155 | 90% | 0% | 85% | 18% | 11% |
| Bulgaria* | 236 | 236 | 25% |  | 93% | 7% | 8% |
| Turkey | 16 | 31 | 94% | 31% | 19% | 0% | 0% |

*\*Interviews in Bulgaria and Greece were conducted in 2012 and 2013 and no* *information on habitat availability was obtained.*

Given that predation by carnivores was the most frequently stated problem, many of the solutions provided revolved around the topic how livestock could be protected from predators, except in Turkey, where rural stakeholders accepted the loss of livestock to natural predators. Between 0 – 93% of respondents mentioned that they use guarding dogs, fenced enclosures or other non-lethal protection measures to keep their livestock safe. The most common alternative solutions to control the problem of predators were shooting all predators (in Syria, Saudi Arabia, and Ethiopia), or the use of poison, which occurred in every country where we interviewed people except in Turkey. The highest prevalence of poison use occurred in Syria and Greece, followed by Saudi Arabia, Egypt, North Macedonia, and Ethiopia (Table 3). Poisoning was rarely admitted in Bulgaria and Albania. However, supplementary data obtained from more detailed and directed conversations with shepherds in Egyptian Vulture territories in Albania confirmed that only 5% of shepherds admitted to using poison themselves, while 43% stated that other shepherds in the area used poison to kill wolves after an attack on their sheep. Similarly, in Greece 63% of stakeholders stated that poisoning occurred, while only 1.1% admitted to use poison themselves. Reporting this widespread but illegal practice is therefore highly variable among stakeholders and countries.

Agricultural chemicals (e.g. strychnine, methomyl) to control feral dogs were used by local authorities in Ethiopia, Egypt, Lebanon, Syria, and Saudi Arabia, and the disposal of poisoned animals was not always safe. Mass poisoning campaigns were conducted for public health reasons in the vicinity of human settlements and at rubbish dumps. Feral dogs poisoned with strychnine or other toxins were mostly buried, but sometimes also disposed in open fields or landfills, which allowed scavenging birds and other carnivores to access and feed on poisoned carcasses. One of our tracked Egyptian Vultures was poisoned on a rubbish dump in Turkey in May 2020. The widespread use of chemical pesticides to suppress locust swarms in eastern Africa and Saudi Arabia began after heavy rains led to exceptionally large locust swarms at the end of 2019 (Roussi 2020). In Saudi Arabia the unresolved loss of one satellite-tracked Egyptian Vulture in April 2020 coincided with the aerial application of chemical pesticides to suppress locusts.

We found that veterinary medical products that are known to cause liver failure in vultures (Diclofenac, Ketoprofen) were widely available in veterinary pharmacies in Egypt, Saudi Arabia, Syria, Jordan, and Lebanon, and available to authorised users in Nigeria. Animals treated with such products were mostly disposed openly in fields or in landfills. On breeding grounds in Greece, veterinarians and livestock holders relied primarily on Meloxicam (which is safe for vultures) and Flunixin (which is toxic to vultures; (Cuthbert *et al*. 2007, Eleni *et al*. 2019)) to treat moribund livestock, and also disposed of carcasses of treated animals openly in fields. In Bulgaria, Diclofenac destined for human use (e.g. standard painkillers) or imported from neighbouring countries was used by small-scale livestock holders without veterinary guidance.

### Direct persecution

Illegal killing of migratory birds in the nine countries we assessed exceeded 6 million birds every year, and may kill >20 million birds in these focal countries alone (Table 4). Because Egyptian Vultures are relatively rare, and do not appear to be targeted specifically even in countries where raptor persecution is a common pastime (e.g. Saudi Arabia), illegal killing may remove 10 – 90 individuals per year (Table 4). Two of our satellite-tracked birds were shot in Lebanon and in Kurdistan (Fig. 1), known hotspots for illegal persecution (Brochet *et al*. 2019a).

**Table 4.** Summary of the number of birds illegally killed every year in nine focal countries along the flyway. The range of birds killed (minimum – maximum) was assessed based on literature and direct surveys (for more details see Brochet *et al*. 2016, Brochet *et al*. 2019a). Countries are ordered by the number of birds killed every year. No data were available for North Macedonia, Ethiopia, Niger and Nigeria.

| Country | N of illegal killing concentration areas | N of killed birds per year (in thousands) | N of killed Egyptian Vultures per year |
| --- | --- | --- | --- |
| Syria | 10 | 2,882 - 4,875 | 5 - 10 |
| Egypt | 9 | 302 - 10,607 | 1 - 10 |
| Lebanon | 5 | 1,678 - 3,480 | 2 - 4 |
| Saudi Arabia | 0 | 625 - 2,500 | unknown |
| Greece | 8 | 485 - 921 | 0 - 3 |
| Albania | 5 | 206 - 325 | 0 - 3 |
| Turkey | 10 | 25 - 117 | 0 - 50 |
| Bulgaria | 5 | 12 - 64 | 0 - 3 |
| Jordan | 5 | 13 - 22 | 0 - 5 |

One particular aspect of the illegal taking of birds is the trade of vulture products in Africa. We surveyed 12 markets with 25,980 stalls or sellers in Nigeria, and 33 markets with 26,955 stalls or sellers in Niger. In Nigeria, 397 sellers (1.5%) offered vulture products for sale, while in Niger only 63 sellers (0.23%) offered vulture products. Although no Egyptian Vultures were available at the time of the surveys in Nigeria, all sellers (100%) stated that they would sell Egyptian Vultures if their suppliers would deliver them. In Niger, 12 sellers (19% of those offering any vulture product) offered Egyptian Vulture parts (Table 5).

**Table 5.** Summary of vulture parts found or reported to have been sold during market surveys in Nigeria and Niger between 2018 and 2020. No markets were surveyed in other countries because vulture products are not generally available for sale in our other focal countries.

| Species | Niger | Nigeria |
| --- | --- | --- |
| Egyptian Vulture | 22 | 0 |
| Hooded Vulture | 74 | 1595 |
| Lappet-faced Vulture | 5 | 10 |
| Rüppell's Vulture | 5 | 0 |
| White-backed Vulture | 0 | 3 |
| unknown species | 6 | 361 |

Egyptian Vulture parts were unavailable on markets in Nigeria because the species has become rare. According to the sellers themselves, the main reason why the species had disappeared was due to direct persecution for trade. Due to the absence of vultures in Nigeria, sellers in Nigeria obtained only a fraction of vultures they sold from within Nigeria, but sourced their products from Niger, Chad, Cameroon, Burkina Faso, Mali, Senegal, and the Central African Republic. Two of our satellite-tracked birds were shot in Niger, one by a hunter supplying markets in Nigeria (Kret *et al*. 2018), and one to supply a local bushmeat market.

### Synthesis and ranking of threats

Using the quantitative evidence provided above, and the regional knowledge of conservation experts working in each country, we ranked the relative importance of threats for each of our focal countries (Table 6). In Europe, poisoning was the most important threat affecting Egyptian Vultures, while in the Middle East direct persecution (Syria, Lebanon), electrocution (Jordan), and poisoning (Saudi Arabia) were the most important threats (Table 6). In Africa, persecution for the vulture trade was the primary threat in Niger and Nigeria, while electrocution was the key threat in Egypt and Ethiopia (Table 6).

**Table 6.** Summary of the importance of three major anthropogenic threats to Egyptian Vultures along the eastern Mediterranean flyway in 2020. Importance of threats was scored qualitatively from 0 (threat absent or irrelevant) to 5 (ubiquitous and severe), with colour coding for visual emphasis. Note that the threat ranking is intended to prioritise work at a country-level, and while threats may be consistently scored within continents, threat rankings between continents may not be comparable.

| Country |  | Poisoning | Electrocution / Collision | Direct persecution |
| --- | --- | --- | --- | --- |
| Europe | Albania | 4 | 3 | 1 |
|  | North Macedonia | 4 | 2 | 1 |
|  | Bulgaria | 4 | 3 | 2 |
|  | Greece | 5 | 3 | 1 |
| Middle East | Turkey | 2 | 5 | 2 |
|  | Syria | 3 | 2 | 4 |
|  | Lebanon | 3 | 2 | 4 |
|  | Jordan | 2 | 4 | 1 |
|  | Saudi Arabia | 5 | 4 | 2 |
| Africa | Egypt | 3 | 4 | 2 |
|  | Ethiopia | 4 | 5 | 0 |
|  | Nigeria | 2 | 1 | 5 |
|  | Niger | 2 | 1 | 4 |

## Discussion

The key threats to Egyptian Vultures along the Eastern Mediterranean flyway vary among the 13 countries that we included in our assessment, with poisoning and electrocution being the threats that occur in every country along the flyway. Collision and electrocution are major problems in Turkey, Ethiopia, Saudi Arabia, Egypt, and Greece, and direct persecution was a key threat in Syria, Lebanon, Egypt, Niger and Nigeria. These threats will affect Egyptian Vultures, but also hundreds of thousands of storks, buzzards, kites and eagles that use the same flyway (Shirihai *et al*. 2000, Megalli & Hilgerloh 2013, Fülöp *et al*. 2014, Oppel *et al*. 2014), and resident raptors, storks and vultures that share the same habitats in Africa (Santangeli *et al*. 2019a, Buechley *et al*. 2021). Although our work cannot estimate which of the threats has the greatest demographic impact on Egyptian Vultures, our overview of the relative importance of threats allows governments and conservation organisations to prioritise efforts to reduce the most important threats in each country that will affect Egyptian Vultures and many other similar species. Rapid action to mitigate threats can protect migratory birds and thus honour each country’s commitments under the Convention of Migratory Species.

The main weakness of our study is that we cannot quantify the impact of any of the threats we identified on vulture populations, and we cannot quantitatively compare the impact of the different threats on populations. While we found evidence for electrocution of vultures in Ethiopia, it is impossible to translate the number of victims found per km of powerline into a typical demographic parameter such as annual survival probability that would allow a quantitative extrapolation of the impact on populations, especially since our surveys did not account for detection probability and may therefore underestimate the true scale of mortality (Bellan *et al*. 2013, Etterson 2013, Korner-Nievergelt *et al*. 2015). Likewise, while we also found evidence that poisoning occurs in Ethiopia, and two of our satellite-tracked birds were poisoned in Ethiopia, the risk of poisoning and electrocution is measured in different currencies in our approach, and cannot be objectively compared. To overcome this issue, we assembled vulture and conservation experts from each country, and discussed the relative importance of each threat based on our evidence and existing detailed country-level information (Table 6). Nonetheless, there is no guarantee that mitigating the most important threats we identified will immediately revert population declines, especially since other threats such as habitat loss or food shortage may impose further diffuse effects on populations.

The threats we evaluate have been previously known to cause bird mortality at large spatial scales, especially for vultures (Kirby et al. 2008; Ogada et al. 2015; Botha et al. 2017), and our work was not designed to identify previously unknown threats. However, given the number and diversity of threats along the flyway, we provide important guidance about which threats should be addressed first in which geographic region (Table 6). Together with work that has identified the geographic areas where vultures reach highest density and diversity (Buechley *et al*. 2018, Santangeli *et al*. 2019a), there is now sufficient information for governments and international funding agencies to work towards reducing threats to vultures (Safford *et al*. 2019) and migratory birds in general. Mitigating certain threats may require greater effort and better governance than is available in several countries, and we caution that the potential effectiveness of any mitigation solutions should be considered given the socio-political realities in each country (Amano *et al*. 2017, Baynham-Herd *et al*. 2018, Santangeli *et al*. 2019b).

The highest priority for immediate action and the highest potential for immediate benefit is to avoid increasing the level of threat posed by electrocution and collision along the flyway. We found that electrocution and collision risk occurs in every country, and this risk is potentially increasing in many countries due to the expansion of electricity distribution and power generation networks (Flade 2012, Kiesecker *et al*. 2019). Numerous guidelines exist on how to design and deploy bird-safe electricity infrastructure (AEWA 2012, BirdLife International 2015, Martín *et al*. 2019), and our assessments indicate that in some countries such infrastructure is in fact well designed: for example, in Syria most of the low-voltage power distribution networks use a safe pylon design, which resulted in low rates of electrocution and collision (Table 1), while in Jordan all wind power installations require a shut-down on demand system that considerably reduces the collision risk to migratory birds (Khoury 2017, Tomé *et al*. 2017). Temporary shut-downs of wind turbines, as well as bird electrocution at power distribution infrastructure that causes power disruptions, incur substantial economic costs (D’Amico *et al*. 2018, Moreira *et al*. 2018). Safer location of large scale power infrastructure developments based on existing knowledge about bird sensitivities (Loss 2016, Smith & Dwyer 2016, Thaxter *et al*. 2017), and safer designs of transport infrastructure will therefore result in both biodiversity and economic benefits and improved quality of power supply. We therefore urge funders and policy makers to revise the design requirements for power infrastructure and develop infrastructural standards that eliminate the electrocution and collision risk to large migratory birds.

Existing infrastructure that poses high electrocution and collision risk needs to be safeguarded with insulation, visual markers, or other effective mitigation devices to reduce bird mortality (López-López *et al*. 2011, AEWA 2012, Badia-Boher *et al*. 2019, Sánchez *et al*. 2020). Such action is exceptionally urgent in Turkey, Ethiopia, and Saudi Arabia, where we found high levels of bird mortality (Table 1), and where major concentrations occur during migration (Oppel *et al*. 2014, Buechley *et al*. 2018) and in winter (Arkumarev *et al*. 2014, Keijmel *et al*. 2020, Shobrak *et al*. 2020). A previous example from Sudan - where a hazardous powerline (Angelov *et al*. 2013) was replaced with bird-safe infrastructure in 2014 to reduce bird electrocution - suggests that this is practically feasible along the flyway. We therefore believe that safeguarding of existing energy infrastructure will be technologically and financially feasible in other countries along the flyway if policy makers and funders honour their commitments under the Convention of Migratory Species.

Besides electrocution and collision, poisoning was the most widespread threat along the flyway. None of our interviews revealed any intention of people or authorities to deliberately poison vultures or other large birds, and poisoning was therefore invariably an inadvertent consequence of securing personal property and income sources, protect game animals, or attempts to improve public health by controlling the number of feral animals. Solutions to reduce the incidence of poisoning are therefore complicated and require an increase of awareness and law enforcement (Ogada 2014, Richards *et al*. 2018), as well as a diversification of household income sources to reduce total dependency on livestock (Davies & Bennett 2007, Romañach *et al*. 2007, Tsegaye *et al*. 2013). Potential improvements to protect livestock from wild predators are enclosures (Lichtenfeld *et al*. 2015) and guarding dogs (van Bommel & Johnson 2012, McManus *et al*. 2015), and helping rural livestock holders to improve their protection measures may reduce the propensity to use poison. Alternatively, improvements to veterinary care and access to water and food resources may increase the health of livestock populations and could either directly reduce predation risk (Khorozyan *et al*. 2015), or reduce the economic impact of livestock predation if predation losses are offset by other sources of income (Jackson & Wangchuk 2004, Romañach *et al*. 2007). The ultimate cause of livestock predation by natural predators, however, is the loss of natural habitats and wildlife populations that serve as principal prey for predators (Kebede *et al*. 2012, Amaja *et al*. 2016). Large scale restoration of natural habitat and wildlife populations may therefore be one of the best mitigation strategies against livestock predation (Georgiadis *et al*. 2007).

Public health officials aiming to control the populations of feral animals should consider the inadvertent risks to wildlife and human health when using non-selective methods such as toxic chemicals that can affect non-target species. In Ethiopia, for example, staff administering the strychnine for feral dog control were negatively affected by the substance. Strychnine and other highly toxic substances should be banned for general use and removed from the market. Alternative approaches to improve public health by controlling or vaccinating feral animals could combine fertility control and more humane and more targeted removal methods (Sillero-Zubiri & Switzer 2004, Massei *et al*. 2010). Similar bans should be considered for the veterinary use of drugs that are known to be harmful for raptors, such as Diclofenac and Ketoprofen. Following catastrophic declines of vulture populations, India, Nepal and Pakistan banned the veterinary use of Diclofenac in 2006, and we recommend that all countries along flyways used by vultures and other large raptors implement a similar ban to prevent avoidable mortality.

Illegal killing of birds causes the death of >20 million individuals along the Eastern Mediterranean flyway every year (Brochet *et al*. 2016), but our assessment revealed that the indiscriminate killing of all migratory birds in the Middle East likely only causes few (<50) Egyptian Vulture deaths every year in our focal countries. The only known place where Egyptian Vultures are targeted is at Lake Nasser in Egypt, where foreign hunting parties shoot large numbers of raptors, ibises, storks, pelicans and waterfowl for taxidermy. However, direct persecution is also widespread in neighbouring countries (Iraq, Iran, and some of the Gulf States), where an estimated 3000 raptors are taken annually including about 100 Egyptian Vultures mainly from Iraq and Yemen (Brochet *et al*. 2019a). There is considerable uncertainty in our assessments because they rely on interviews and extrapolations, and the true extent of Egyptian Vulture mortality may be higher. More effective legislation and more robust enforcement of existing hunting laws are urgently required in several countries along the flyway.

The direct persecution for the market trade in Nigeria is highly targeted and has the potential to affect all vulture populations in the central Sahel region (Saidu & Buij 2013, Ogada *et al*. 2015, Buij *et al*. 2016), especially neighbouring countries such as Niger. We found hundreds of market sellers willing to sell vulture products, including Egyptian Vultures, and these products may be sourced from other countries (Kret *et al*. 2018), and possibly as a by-product of poisoning incidents (Mateo-Tomás & López-Bao 2020). The trade in vulture parts also exists in Niger, but at a smaller scale, and with much of the demand driven by Nigeria. Vultures are legally protected in Nigeria, with both the killing and trade of vulture species being illegal. However, this law is not enforced and without a reduction in demand in Nigeria, it is unlikely that the persecution in the Sahel region will cease. Reducing the demand in wildlife products is a complicated issue with little evidence for the effectiveness of campaigns so far (Veríssimo & Wan 2019). Clear communication about effectiveness and causality may overcome demand for products that are believed to have health benefits (MacFarlane *et al*. 2018), and efforts are under way in Nigeria to replace vulture products with plant-based alternatives. We encourage governments to enforce existing wildlife laws and reduce the trade in wildlife that has negative effects on biodiversity and human welfare, potentially using the COVID-19 pandemic in 2020 as an example in awareness campaigns (Volpato *et al*. 2020, Zhou *et al*. 2020).

In summary, governments along the Eastern Mediterranean flyway face considerable challenges to protect migratory birds under the Convention of Migratory Species, and the most important actions to reduce mortality will vary by country. While governments on eastern European breeding grounds urgently need to address the problem of wildlife poisoning, the governments in Turkey, Saudi Arabia, and Ethiopia should focus on reducing threats to migratory birds from energy infrastructure while not neglecting the threat of poisoning. Enforcement of existing laws to reduce the illegal killing of birds should be a priority in Syria, Lebanon, Egypt, Nigeria and Niger.

## Acknowledgements

This work was carried out in the framework of the LIFE projects “The Return of the Neophron” (LIFE10 NAT/BG/000152) and “Egyptian Vulture New LIFE” (LIFE16 NAT/BG/000874, www.LifeNeophron.eu) funded by the European Union and co-funded by the A. G. Leventis Foundation and the BirdLife GEF/UNDP Migratory Soaring Birds project. We appreciate the support during fieldwork by Lubomir Peske, Clementine Bougain, Mekonnen Kassa, Samson Zelleke, Behailu Abraham, Nora Juhar, and Amin Seid. Permission to conduct research in Ethiopia was granted by the Ethiopian Wildlife Conservation Authority and we appreciate the support of Dr Fanuel Kebede. We thank Joelene Hughes, Sorrel Jones, Michael McDonald, Richard Bradbury, Andrea Santangeli and Vanessa Berrie for helpful discussions how to design questionnaires and evaluate responses, and Juliet Vickery for constructive comments on a previous draft of the manuscript. We appreciate the support of The WindPower.net, represented by Michael Pierrot, for supplying proprietary information about the location and size of wind power installations.

## Supplementary Information

**Table S1:** Complete summary of interviews with rural stakeholders to understand common problems with land use and livestock herding, and solutions adopted to overcome these problems. Note that poisoning is referred to at a community rather than at an individual level, and its prevalence may therefore not be comparable to shooting or protection in some countries; ‘Habitat / Grazing land’ refers to the problem of insufficient grazing or crop land.

| Country | Response | Category | Proportion respondents |
| --- | --- | --- | --- |
| Albania | Problem | Predation | 0.899 |
| Albania | Problem | Other | 0.101 |
| Albania | Problem | Drought / Climate | 0.000 |
| Albania | Problem | Disease | 0.000 |
| Albania | Problem | Invasive species | 0.000 |
| Albania | Problem | Habitat / Grazing land | 0.000 |
| Albania | Solution | Guard/Chase | 0.851 |
| Albania | Solution | Shooting | 0.182 |
| Albania | Solution | Poison | 0.108 |
| Albania | Solution | Other | 0.101 |
| Albania | Solution | Trapping | 0.014 |
| Albania | Solution | Fencing | 0.000 |
| Albania | Solution | Medicine | 0.000 |
| Albania | Solution | Migration | 0.000 |
| Bulgaria | Problem | Predation | 0.254 |
| Bulgaria | Problem | Drought / Climate | 0.059 |
| Bulgaria | Problem | Disease | 0.004 |
| Bulgaria | Problem | Invasive species | 0.000 |
| Bulgaria | Problem | Habitat / Grazing land | 0.000 |
| Bulgaria | Solution | Predation | 0.254 |
| Bulgaria | Solution | Guard/Chase | 0.106 |
| Bulgaria | Solution | Poison | 0.076 |
| Bulgaria | Solution | Shooting | 0.008 |
| Bulgaria | Solution | Fencing | 0.004 |
| Bulgaria | Solution | Trapping | 0.000 |
| Bulgaria | Solution | Other | 0.000 |
| Bulgaria | Solution | Medicine | 0.000 |
| Bulgaria | Solution | Migration | 0.000 |
| Egypt | Problem | Drought / Climate | 0.500 |
| Egypt | Problem | Predation | 0.500 |
| Egypt | Problem | Disease | 0.250 |
| Egypt | Problem | Habitat / Grazing land | 0.250 |
| Egypt | Problem | Invasive species | 0.000 |
| Egypt | Problem | Other | 0.000 |
| Egypt | Solution | Other | 0.750 |
| Egypt | Solution | Poison | 0.500 |
| Egypt | Solution | Guard/Chase | 0.250 |
| Egypt | Solution | Shooting | 0.000 |
| Egypt | Solution | Trapping | 0.000 |
| Egypt | Solution | Fencing | 0.000 |
| Egypt | Solution | Medicine | 0.000 |
| Egypt | Solution | Migration | 0.000 |
| Ethiopia | Problem | Predation | 0.868 |
| Ethiopia | Problem | Drought / Climate | 0.426 |
| Ethiopia | Problem | Disease | 0.382 |
| Ethiopia | Problem | Invasive species | 0.368 |
| Ethiopia | Problem | Habitat / Grazing land | 0.294 |
| Ethiopia | Problem | Other | 0.074 |
| Ethiopia | Solution | Shooting | 0.676 |
| Ethiopia | Solution | Poison | 0.456 |
| Ethiopia | Solution | Fencing | 0.265 |
| Ethiopia | Solution | Medicine | 0.118 |
| Ethiopia | Solution | Other | 0.118 |
| Ethiopia | Solution | Guard/Chase | 0.088 |
| Ethiopia | Solution | Migration | 0.088 |
| Ethiopia | Solution | Trapping | 0.059 |
| Greece | Problem | Predation | 0.469 |
| Greece | Problem | Drought / Climate | 0.095 |
| Greece | Problem | Disease | 0.014 |
| Greece | Problem | Invasive species | 0.000 |
| Greece | Problem | Habitat / Grazing land | 0.000 |
| Greece | Solution | Poison | 0.627 |
| Greece | Solution | Predation | 0.469 |
| Greece | Solution | Guard/Chase | 0.226 |
| Greece | Solution | Shooting | 0.076 |
| Greece | Solution | Other | 0.065 |
| Greece | Solution | Fencing | 0.054 |
| Greece | Solution | Trapping | 0.005 |
| Greece | Solution | Medicine | 0.000 |
| Greece | Solution | Migration | 0.000 |
| Lebanon | Problem | Habitat / Grazing land | 0.455 |
| Lebanon | Problem | Disease | 0.318 |
| Lebanon | Problem | Predation | 0.227 |
| Lebanon | Problem | Other | 0.045 |
| Lebanon | Problem | Drought / Climate | 0.000 |
| Lebanon | Problem | Invasive species | 0.000 |
| Lebanon | Solution | Shooting | 0.318 |
| Lebanon | Solution | Poison | 0.273 |
| Lebanon | Solution | Medicine | 0.273 |
| Lebanon | Solution | Fencing | 0.136 |
| Lebanon | Solution | Other | 0.045 |
| Lebanon | Solution | Trapping | 0.000 |
| Lebanon | Solution | Guard/Chase | 0.000 |
| Lebanon | Solution | Migration | 0.000 |
| Niger | Problem | Predation | 0.667 |
| Niger | Problem | Habitat / Grazing land | 0.500 |
| Niger | Problem | Drought / Climate | 0.167 |
| Niger | Problem | Disease | 0.167 |
| Niger | Problem | Invasive species | 0.000 |
| Niger | Problem | Other | 0.000 |
| Niger | Solution | Guard/Chase | 0.667 |
| Niger | Solution | Poison | 0.333 |
| Niger | Solution | Fencing | 0.167 |
| Niger | Solution | Shooting | 0.000 |
| Niger | Solution | Trapping | 0.000 |
| Niger | Solution | Medicine | 0.000 |
| Niger | Solution | Migration | 0.000 |
| Niger | Solution | Other | 0.000 |
| North Macedonia | Problem | Predation | 0.419 |
| North Macedonia | Problem | Habitat / Grazing land | 0.323 |
| North Macedonia | Problem | Other | 0.161 |
| North Macedonia | Problem | Drought / Climate | 0.097 |
| North Macedonia | Problem | Disease | 0.000 |
| North Macedonia | Problem | Invasive species | 0.000 |
| North Macedonia | Solution | Guard/Chase | 0.774 |
| North Macedonia | Solution | Poison | 0.484 |
| North Macedonia | Solution | Shooting | 0.194 |
| North Macedonia | Solution | Other | 0.194 |
| North Macedonia | Solution | Trapping | 0.000 |
| North Macedonia | Solution | Fencing | 0.000 |
| North Macedonia | Solution | Medicine | 0.000 |
| North Macedonia | Solution | Migration | 0.000 |
| Syria | Problem | Predation | 1.000 |
| Syria | Problem | Habitat / Grazing land | 1.000 |
| Syria | Problem | Drought / Climate | 0.250 |
| Syria | Problem | Disease | 0.000 |
| Syria | Problem | Invasive species | 0.000 |
| Syria | Problem | Other | 0.000 |
| Syria | Solution | Shooting | 1.000 |
| Syria | Solution | Poison | 1.000 |
| Syria | Solution | Trapping | 0.000 |
| Syria | Solution | Guard/Chase | 0.000 |
| Syria | Solution | Fencing | 0.000 |
| Syria | Solution | Medicine | 0.000 |
| Syria | Solution | Migration | 0.000 |
| Syria | Solution | Other | 0.000 |
| Turkey | Problem | Predation | 0.938 |
| Turkey | Problem | Drought / Climate | 0.875 |
| Turkey | Problem | Disease | 0.812 |
| Turkey | Problem | Habitat / Grazing land | 0.312 |
| Turkey | Problem | Invasive species | 0.000 |
| Turkey | Problem | Other | 0.000 |
| Turkey | Solution | Medicine | 0.938 |
| Turkey | Solution | Guard/Chase | 0.188 |
| Turkey | Solution | Shooting | 0.000 |
| Turkey | Solution | Poison | 0.000 |
| Turkey | Solution | Trapping | 0.000 |
| Turkey | Solution | Fencing | 0.000 |
| Turkey | Solution | Migration | 0.000 |
| Turkey | Solution | Other | 0.000 |

## Notes

https://www.movebank.org/cms/webapp?gwt_fragment=page=studies,path=study15869951

### Competing Interest Statement

The authors have declared no competing interest.

